# Time-resolved analyses of elemental distribution and concentration in living plants: An example using manganese toxicity in cowpea leaves

**DOI:** 10.1101/228577

**Authors:** F. Pax C. Blamey, David J. Paterson, Adam Walsh, Nader Afshar, Brigid A. McKenna, Miaomiao Cheng, Caixan Tang, Walter J. Horst, Neal W. Menzies, Peter M. Kopittke

## Abstract

- Knowledge of elemental distribution and concentration within plant tissues is crucial in the understanding of almost every process that occurs within plants. However, analytical limitations have hindered the microscopic determination of changes over time in the location and concentration of nutrients and contaminants in living plant tissues.
- We developed a novel method using synchrotron-based micro X-ray fluorescence (μ-XRF) that allows for laterally-resolved, multi-element, kinetic analyses of plant leaf tissues *in vivo*. To test the utility of this approach, we examined changes in the accumulation of Mn in unifoliate leaves of 7-d-old cowpea (*Vigna unguiculata*) plants grown for 48 h at 0.2 and 30 μM Mn in solution.
- Repeated μ-XRF scanning did not damage leaf tissues demonstrating the validity of the method. Exposure to 30 μM Mn for 48 h increased the initial number of small spots of localized high Mn and their concentration rose from 40 to 670 mg Mn kg^-1^ fresh mass. Extension of the two-dimensional μ-XRF scans to a three-dimensional geometry provided further assessment of Mn localization and concentration.
- This method shows the value of synchrotron-based μ-XRF analyses for time-resolved *in vivo* analysis of elemental dynamics in plant sciences.

## Introduction

The concentration and distribution of nutrients and contaminants within plant tissues change over time in response to physiological stimuli, developmental stage, and changes in the broader external environment. To understand the underlying genetic and physiological processes influencing plant growth, it is necessary to determine the concomitant changes in the accumulation or decline of these elements within plant tissues. In this regard, ionomics is concerned with the examination of elements in plants, although measurements are normally conducted for bulk tissues (Salt *et al.*, 2008). Whilst such analyses provide valuable information, it is even more useful to determine laterally resolved concentrations of essential and non-essential elements within plant tissues (Conn & Gilliham, 2010). Besides its relevance in plant nutrition, ionomics involves the functional analysis of genes that directly and indirectly control plant development and physiology (Salt *et al*., 2008; Takahashi *et al*., 2009).

Various techniques exist for examining the distribution of elements within plant tissues but most techniques require extensive processing of sequential samples. Hence, they are not able to examine nutrient and contaminant changes over time in the same area of living plants. For example, conventional scanning electron microscopy coupled with energy-dispersive X-ray spectroscopy (SEM-EDS) needs an ultra-high vacuum that requires samples to be dehydrated; frozen samples may be used where a cryo-SEM-EDS is available (Cosio *et al.*, 2005). It is perhaps possible to examine living plants using environmental SEM (ESEM), but there are problems with sample size restrictions, electron beam damage, and a comparatively poor detection limit (Danilatos, 1981; McGregor & Donald, 2010). Though having excellent subcellular resolution, nanoscale secondary ion mass spectrometry (NanoSIMS) analysis also requires ultra-high vacuum (Moore *et al.*, 2014). Confocal microscopy with fluorophores is potentially of use for the kinetic analyses of living plants (Walczysko *et al.*, 2000; Babourina & Rengel, 2009) but there are limits imposed by the availability of fluorophores and uncertainty in their selectivity and cellular penetration. Laser ablation inductively coupled plasma mass spectrometry (LA-ICP-MS) is also capable of examining living plants in ambient conditions (Salt *et al.*, 2008) but damage through ablation of the sample surface prevents kinetic analysis of the same tissue area. Autoradiography has been used for the study of plants since the 1920s (Hevesy, 1923), and macro-autoradiography can potentially be used for kinetic analyses of living plants but is limited by poor resolution, long exposure times, limited availability of suitable isotopes, and safety considerations (Solon *et al.*, 2010). Recently-developed radioisotope tracer techniques for *in vitro* analysis, such as a positron-emitting tracer imaging system (Tsukamoto *et al*., 2006) and magnetic resonance imaging (Jahnke *et al*., 2009), have overcome some limitations, but it is only possible to examine a single element at a time (Sugita *et al*., 2016).

Synchrotron-based micro-X-ray fluorescence spectroscopy (μ-XRF) is of interest as there are no theoretical restrictions on sample size, analyses are conducted at ambient temperature and pressure often at a resolution of *c.* ≤ 1 μm with a detection limit of *c*. 0.1 to 100 mg kg^-1^ fresh mass (FM). Leaf tissues of living plants may be examined also (Scheckel *et al.*, 2004). Depending upon the element of interest and beamline specifications, this technique can simultaneously generate maps for multiple elements within the energy range of the beamline, often 2 to 25 keV allowing analysis from P to Ag at the K-edge. Early analyses of plant tissues using μ-XRF analyses focused on dehydrated tissues of hyperaccumulators (McNear *et al.*, 2005; McNear & Küpper, 2014). However, tissue dehydration may result in experimental artifacts and the high concentrations used in studies of hyperaccumulators are not relevant to many crop species. With progressive improvements in technology (specifically, the development of more efficient fluorescent X-ray detectors), there has been increasing interest in the analysis of hydrated tissues of non-hyperaccumulating species (Lombi *et al.*, 2011a; Blamey *et al.*, 2015; Kopittke *et al.*, 2015). For example, studies of Cu, Ni, Zn, and Mn rhizotoxicity in hydrated cowpea (*Vigna unguiculata*) roots showed Cu located in the rhizodermis and outer cortex, Ni in the inner cortex, and Zn in the stele; the meristematic zone was high in both Zn and Mn (Kopittke *et al.*, 2011; Kopittke *et al.*, 2013). However, problems may still arise with radiation damage and leaf tissue dehydration by high-energy X-rays upon repeated scanning of the same area of leaf (Lombi & Susini, 2009) that would preclude repeated analysis of living tissues.

To examine the potential of synchrotron-based μ-XRF for the analysis of living plants, we inspected changes in element distribution in leaves of cowpea following exposure to an adequate and a toxic level of Mn in the rooting medium. Using μ-XRF, we previously identified high Mn in sunflower (*Helianthus annuus*) trichomes and in vacuoles of white lupin (*Lupinus albus*) as mechanisms of tolerance (Blamey *et al.*, 2015). Such mechanisms are not present in cowpea (an established model species in Mn toxicity studies) and soybean (*Glycine max*), making these crop species considerably more sensitive to high Mn^2+^ in the root environment that results from the localized accumulation of Mn in leaf tissues (Heenan & Carter, 1976; Horst, 1983). The first visible symptom of toxicity is the appearance of small dark spots on unifoliate leaves *c.* 4-d after exposure of roots to 20 μM Mn (Wissemeier & Horst, 1987). Thus, the present study tested the suitability of μ-XRF for *in situ*, multi-element, kinetic, microscopic analyses to quantify changes in elemental distribution in leaves of living cowpea plants. We envisage that the method developed here will be of importance across a wide range of studies for the examination of plant responses to physiological stimuli, developmental stage, and changes in biotic and abiotic environments.

## Materials and Methods

### Plant growth

Cowpea (*Vigna unguiculata* L. Walp. cv. Bunya) seeds in rolled paper towels were placed in tap water and seedlings transplanted 4 d later into 20 L of aerated nutrient solution at pH 5.6 and ionic strength of *c.* 3 mM (Blamey *et al.*, 2015) approximating that in soil solutions (Kopittke *et al.*, 2010). Nominal concentrations of nutrients in the basal solution were (μM): 1000 Ca, 120 NH_4_^+^-N, 95 Mg, 300 K, 10 Na, 6 Fe, 0.2 Mn, 0.5 Zn, 0.2 Cu, 1250 Cl, 670 NO_3_^-^-N, 340 S, 5 P, 1 B, and 0.01 Mo. After 7 d in a controlled environment room at 25 °C under fluorescent lights, plants were transferred to the Australian Synchrotron and grown in fresh nutrient solutions at 22 °C under high-pressure sodium lights at photosynthetically active radiation (PAR) of 1,500 μmol m^-2^ s^-1^. Mean concentrations of selected nutrients measured by inductively coupled optical emission spectroscopy (ICP-OES) in solutions at the beginning and end of the initial and final periods were (μM): 1,050 Ca, 120 Mg, 360 K, 30 Na, 0.23 Mn, 10 Fe, 0.4 Zn, 0.2 Cu, 360 S, 7 P, and 4 B. Use of 15 % NH_4_^+^-N in solution ensured that the initial solution pH 5.6 did not require adjustment.

A preliminary experiment tested if repeated μ-XRF scans (see below) of the same area of leaf caused damage that would potentially modify the transport of solutes in the xylem and phloem or cause other experimental artifacts (e.g. dehydration). This involved excision of a unifoliate leaf followed by mounting between two layers of 4-μm Ultralene^®^ film on a specimen holder that provided support and limited dehydration (Fig. 1a).

**Fig. 1.**
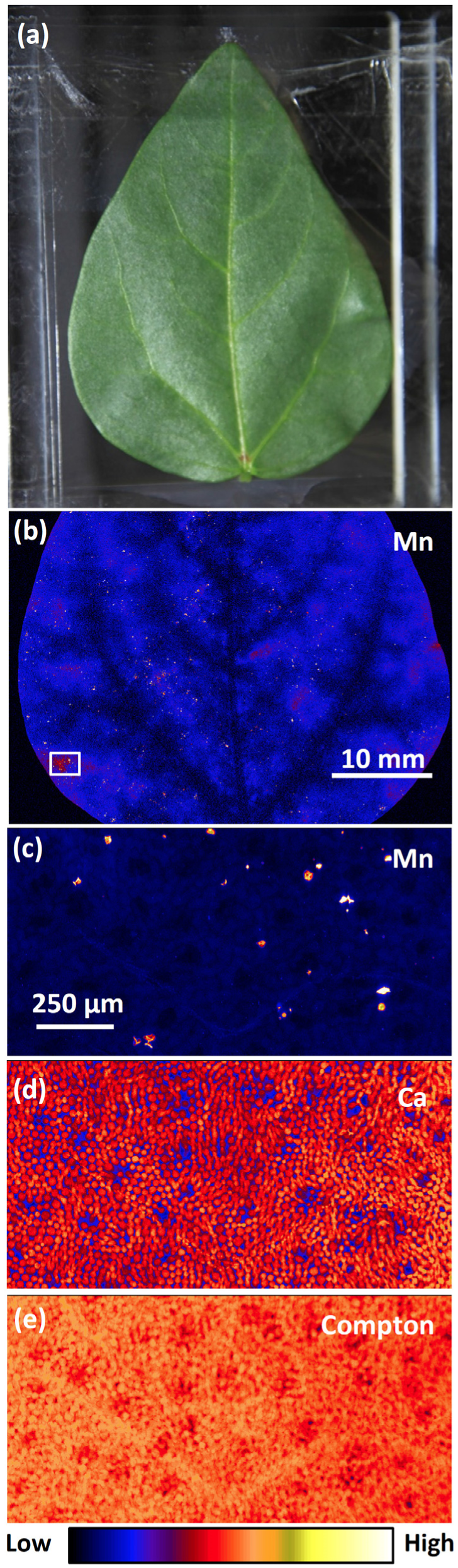
Images of a detached unifoliate leaf of cowpea grown for 7 d at 0.2 μM Mn after conducting a survey scan and three detailed scans. (a) Optical image of a leaf mounted between Ultralene^®^ films. (b) Survey μ-XRF scan of Mn distribution, the white box showing the area of subsequent detailed scans. (c,d) Detailed μ-XRF scan showing the distributions of Mn and Ca. (e) Detailed Compton scatter.

The main experiment involved the transfer of four plants, each to a 50-mL polypropylene centrifuge tube (30 × 115 mm) with roots submerged in 40 mL of basal nutrient solution. Securing the plant stem in the neck of the tube by rolling a length of 15-mm wide geotextile covered by a slotted cap and Parafilm M^®^ prevented spillage onto sensitive equipment. Also secured in the neck of the centrifuge tube were two 2-m lengths of PTFE #2 AWG thin-wall tubing (Cole-Parmer Instrument Company, Vernon Hills IL, USA), one serving as solution input and the other as output. Tygon^®^ tubing with 0.7-mm internal diameter joined the PTFE tubing where necessary. The input tube connected a 10-L reservoir of continuously aerated nutrient solution via a four-roller peristaltic pump to the bottom of the solution in the centrifuge tube. The output tube extended from the 40-mL mark of the centrifuge tube to the reservoir via the peristaltic pump, adjustment of which ensured no solution overspill. Circulation of nutrient solution between the reservoir and the centrifuge tube at a measured rate of 4 mL min^-1^ enabled 90 % renewal of the 40 mL of nutrient solution in < 30 min. Two purpose-built sample holders, each with two centrifuge tubes, contained plants fixed to each holder. Securing a unifoliate leaf of each plant with Ultralene^®^ and double-sided tape protected the sensitive Be detector window *c*. 1 mm away but also allowed transpiration via both the adaxial and abaxial leaf surfaces (Fig. 2a). Each sample holder (i.e. replicate) with two secured plants was attached in turn to the sample stage that allowed horizontal and vertical adjustment of the leaves and correct focusing of the X-ray beam (Fig. 2b). With solution in each centrifuge tube connected to a 10 L-reservoir, we imposed two treatments with nominal 0.2 (i.e. basal) and 30 μM Mn. Mean measured values at the start and end of the 48-h experimental period were 0.33 and 30 μM Mn.

**Fig. 2.**
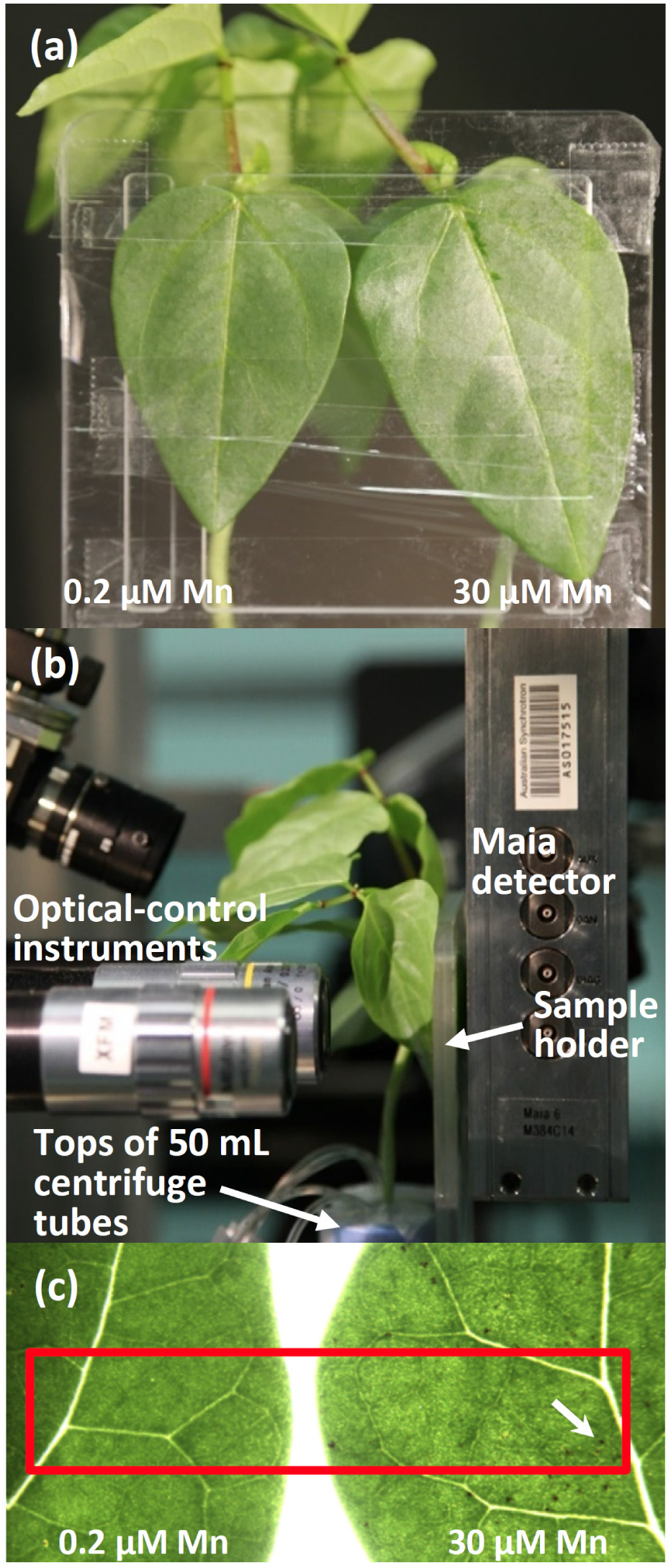
Optical images of cowpea unifoliate leaf arrangement in the sample holder. (a) Leaves of two plants grown at 0.2 and 30 μM Mn between Ultralene^®^ films in the sample holder. (b) Arrangement of living plants mounted in the X-ray beam. (c) Micrograph of unifoliate leaves after the detailed μ-XRF scan at 48 h, the red box showing the area of sequential detailed scans and the white arrow a visible dark spot of Mn accumulation at 30 μM Mn.

An important objective of the main experiment was ensuring optimal plant growth conditions other than in the 30 μM Mn treatment. We had initially intended to renew the 40 mL of solution in each centrifuge tube at appropriate intervals, but an anticipated rapid decrease in dissolved O_2_ precluded this approach. Specifically, we calculated that a 7-d-old root system (2.5 g FM plant^-1^) with an O_2_ consumption rate of 20 μmol g^-1^ FM h^-1^ (Bravo & Uribe, 1981) would deplete the 510 μmol dissolved O_2_ in 40 mL solution at 25 °C in < 30 min. Thus, root O_2_ and nutrient requirements over 48 h were met by circulating 10 L of aerated solution. Use of a peristaltic pump and reservoirs placed on the floor proved satisfactory provided the outflow rate exceeded that of the inflow to prevent overflow of the 50-mL centrifuge tube. This ensured the safety of sensitive electronic equipment that had previously been avoided through use of solid growth medium (Scheckel *et al.*, 2004) or a small reservoir of solution with *Arabidopsis thaliana* plants (Fittschen *et al.*, 2017). Additionally, we had to prevent damage to the fragile Be window of the Maia detector despite the need for unifoliate leaves being at a distance of *c*. 1 mm for correct focus (http://www.synchrotron.org.au/aussyncbeamlines/x-ray-fluorescence-microscopy/samples). Securing the unifoliate leaves to the sample holder with Ultralene^®^ and double-sided tape (Fig. 2a) and moving the other leaves away from the detector (Fig. 2 b) overcame this potential problem.

The final experiment of this study determined the bulk concentration of Mn in unifoliate leaves of cowpea plants grown in a laboratory under high-pressure sodium lights at 25 °C using the same procedure as above. Four plants in each of six 20-L pots were grown in basal nutrient solution for 7 d after transplanting at which time three replicates of 0.2 and 30 μM Mn were imposed for 2 d. Mean concentrations of selected nutrients in solution measured by ICP-OES were (μM): 820 Ca, 85 Mg, 50 K, 20 Na, 10 Fe, 0.28 Zn, 0.12 Cu, 150 S, 10 P, and 2.7 B. The mean concentration of Mn was 0.23 and 28 μM in the two treatments at the start and end of 2 d. Four unifoliate leaves were harvested from each pot, FM and dry mass (DM) determined, and leaves digested in 5:1 HNO_3_:HClO_4_ prior to Mn analysis using ICP-OES.

### μ-XRF scans

Details of the XFM beamline at the Australian Synchrotron have been provided by Paterson *et al.* (2011), as have details for the analysis of fresh hydrated roots by Kopittke *et al.* (2011) and leaves by Blamey *et al.* (2015). X-rays, at an energy of 12.9 keV in the present study, were selected using a Si(111) monochromator and focused (2 × 2 μm) by a pair of Kirkpatrick-Baez mirrors and the X-ray fluorescence emitted by the specimen collected in a backscatter geometry using a 384-element Maia detector system (Lombi *et al.*, 2011a; Paterson *et al.*, 2011). Elemental mapping is conducted on-the-fly in the horizontal direction. Until recently, discrete steps in the vertical direction increased the dwell at the edge of the scanned area (i.e. at the end of each horizontal line scan) with greater potential for X-ray damage to hydrated tissues. A recent improvement allows significantly faster and more accurate raster scanning via kinematically optimized fly-scans known as Rascan, a system that wraps the motion controller into an optimal two-dimension scanner. This device renders a motion trajectory optimized for smoothness, as well as minimal overhead times, for a given fly-scan of known transit time (dwell) and pitch. Importantly, Rascan reduces end-of-line overheads from > 350 ms to *c*. 35 ms resulting in a *c*. 30 % reduction in scan time and less radiation damage. Using this approach, analyses are routinely conducted with a dwell of ≤ 1 ms pixel^-1^, comparing favorably to many other synchrotron-based μ-XRF beamlines where a dwell of 10 to 100 ms pixel^-1^ is common (Lombi *et al.*, 2011b). This decreased dwell using the Maia detector system and Rascan at the Australian Synchrotron facilitated the present study with repeated scanning of the same area of living plant leaf tissues.

The preliminary experiment determined if sequential scanning of the same area resulted in damage from high energy X-rays. The survey scan started within 5 min of excising and mounting a unifoliate leaf (Fig. 1a) and took 52 min to complete an area of 47 × 33 mm with a step size (i.e. virtual pixel size) of 50 μm. From this survey scan, a 7 × 3 mm area was selected for further examination with three detailed scans with a step size of 5 μm, each scan taking *c*. 20 min. Each of the detailed scans covered a slightly larger area encompassing the entire previous scan to test if the X-rays damaged the leaf that would be evident in the subsequent scans by changes in sample hydration and elemental redistribution. A fourth, high-resolution scan with a step size of 2 μm covering an area of 1.54 × 0.79 mm tested the possibility of damage in detail. We also used light microscopy to examine the leaf for any visible signs of damage.

Thereafter, the main experiment involved securing the sample holder with two centrifuge tubes, each with a living plant (Fig. 2), in the beamline and verifying basal solution (0.2 μM Mn) flow rate. Addition of a 0.46-mL aliquot of 0.65 M MnSO_4_ stock solution to one of the 10-L reservoirs imposed a 30-μM Mn treatment. Two initial survey scans of 6 and 2 min identified the area for detailed scans (Fig. 2c), the first of which started immediately thereafter (0 h). This and subsequent detailed scans took *c.* 120 min to complete. There were six scans from 0-2, 6-8, 12-14, 18-20, 24-26, and 48-50 h after first imposition of Mn treatments. The same procedures and scans of adjacent unifoliate leaves followed with living plants in the second replicate. In the replicate reported here, the detailed scan area was *c.* 120 mm^2^, which included a leaf area of 55.2 and 56.4 mm^2^ in the 0.2 and 30 μM Mn treatments. The detailed scans had 4 × 4 μm pixels and a velocity of 4 mm s^-1^ resulting in a pixel transit time of 1 ms. With a total photon flux of *c*. 2 × 10^9^ photon s^-1^ at an energy of 12.9 keV, these scan parameters corresponded to *c*. 1.3 × 10^5^ photon μm^-2^ for each scan (i.e. a total of 8.0 × 10^5^ photon μm^-2^ for the six scans).

On completion of the scans, the CSIRO Dynamic Analysis method in GeoPIXE provided quantitative, true-element images of X-ray fluorescence spectra (Ryan & Jamieson, 1993; Ryan, 2000) as outlined at http://www.nmp.csiro.au/dynamic.html. The survey scans required correction for variation in leaf thickness arising from the major leaf veins by normalizing to Compton scatter. Sections of leaves for detailed scans had only a few minor veins (Fig. 2c) that did not interfere greatly with determination of the two-dimensional areal concentrations of elements and their distributions (Kopittke *et al.*, 2011).

GeoPIXE provided initial quantitative analysis of Mn concentration in 2.6 × 0.08 mm transects of detailed scans across the leaf at 30 μM Mn from 0 to 48 h that involved 650 individual pixels in the horizontal direction × the mean of 20 pixels in the vertical direction. This was followed by determining Mn in the high-resolution image area of 2.6 × 0.6 mm (i.e. values of individual 650 × 150 pixels). Further analysis using GeoPIXE and ImageJ 1.48v (Schneider *et al.*, 2012) extended the two-dimension μ-XRF scans to a three-dimensional geometry of Mn distribution and concentration. The number and localized areas of Mn accumulation and the distributions of K, Ca, Fe, Cu, and Zn at 0 and 48 h in the two Mn treatments were determined also.

## Results

### Potential X-ray damage to living unifoliate leaves

The Australian Synchrotron XFM beamline Maia detector system with fast data acquisition has the advantage of rapid pixel transit times (Paterson *et al.*, 2011) but sequential *in situ* μ-XRF analyses of hydrated leaf tissues may still result in radiation damage and tissue dehydration as shown in micro-tomography of a cowpea root (Lombi *et al.*, 2011a). This was evident also in an unpublished study using X-ray absorption near edge structure (XANES) imaging of a 4.0 × 2.5 mm section of a soybean leaf at high Mn with repeated scans of the same area at increasing energy. Compared to the rest of the leaf, the rectangular scanned area was mildly chlorotic suggestive of radiation damage (Fig. S1a). Furthermore, there were changes in the distribution of Ca (Fig. S1b) and dehydration damage evident in Compton scatter (Fig. S1c) though not in Mn distribution (data not presented). Although mapping was conducted on-the-fly in the horizontal direction with a dwell of 1 ms per virtual 0.1-mm step, damage occurred in a previous study at ends of the horizontal scan lines because of the increased dwell of discrete steps in the vertical direction. In contrast to the effects observed in this previous XANES imaging study, the preliminary experiment in the present study produced no evidence of radiation damage or dehydration to a cowpea leaf after scanning for *c*. 2 h (Fig. 1). This was attributed to the newly-implemented Rascan system which markedly reduced overheads, and hence reduced dwell at the end of the horizontal lines during the vertical move to the next line. We therefore concluded that rapid scanning prevented leaf damage suggesting that experimental artifacts would not arise with time-resolved μ-XRF scans of living leaf tissues.

### Changes in elemental distribution and concentration

There were no visible symptoms of Mn toxicity on leaves of plants grown for 7 d with the basal 0.2 μM Mn in solution (Fig. 1a) but there were a few localized spots of Mn accumulation visible in the μ-XRF scans of the detached unifoliate leaf during the preliminary experiment (Fig. 1b,c). This was evident also at 0 h in both living cowpea unifoliate leaves (Fig. 3) but it was only after 2 d at 30 μM Mn that dark spots indicative of Mn accumulation were visible (Fig. 2c).

**Fig. 3.**
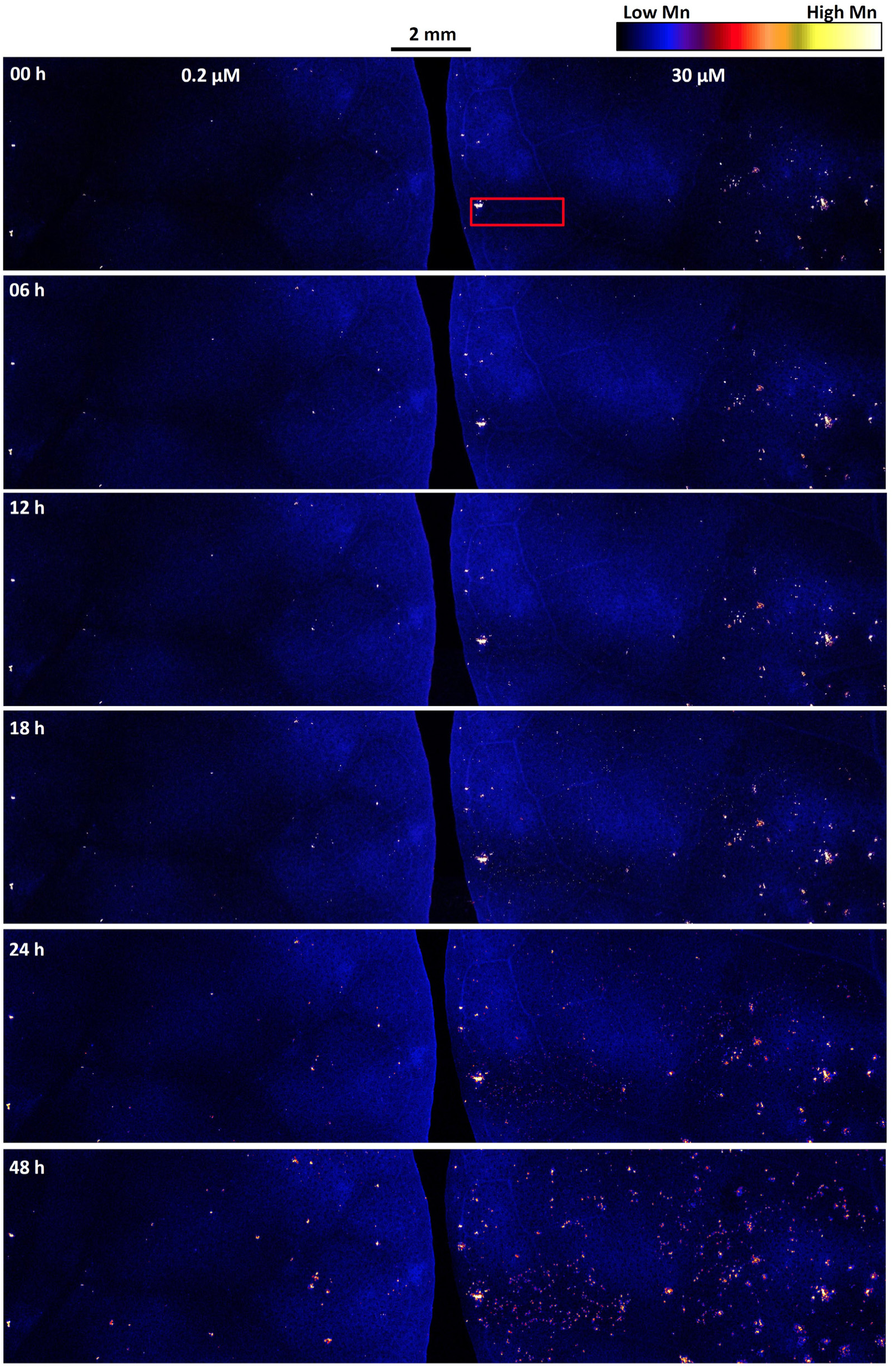
Detailed μ-XRF scans of Mn distribution in unifoliate leaves of cowpea from 0 to 48 h after initial exposure to 30 μM Mn in solution culture. The red box at 0 h shows the area analyzed at high resolution for the distribution of Mn (Fig. 4), for three-dimensional representation of Mn distribution and concentration (Fig. 5), and for ImageJ and GeoPIXE analyses (Fig. 6).

Visual assessment of the detailed μ-XRF scans (Fig. 3) indicated a slight increase over time in the number of high-Mn spots at 0.2 μM Mn, but this increase was considerably lower than that with 30 μM Mn in solution. Given that each detailed scan was *c*. 8.5 megapixels, we selected a small area of the image of 1.56 mm^2^ (2.6 × 0.6 mm) to examine changes in elemental distribution in the 30-μM Mn treatment over the 48-h experimental period. At the start of the experiment, high Mn spots were visible to the left of the image, the largest being *c*. 240 × 130 μm in size (Fig. 4). Elsewhere, Mn was minimally above background. There was little change in Mn distribution after 6 h at 30 μM Mn, with some new high Mn spots evident after 12 h and especially from 18 to 48 h. Transects across the images from 0 to 48 h (Fig. 4) showed no visible increase in the number or concentration of high Mn spots at 6 h, a few instances where new high Mn spots were visible at 12 h. It was only from 18 to 48 h, however, that there were clear increases in spot numbers and in their Mn concentration.

**Fig. 4.**
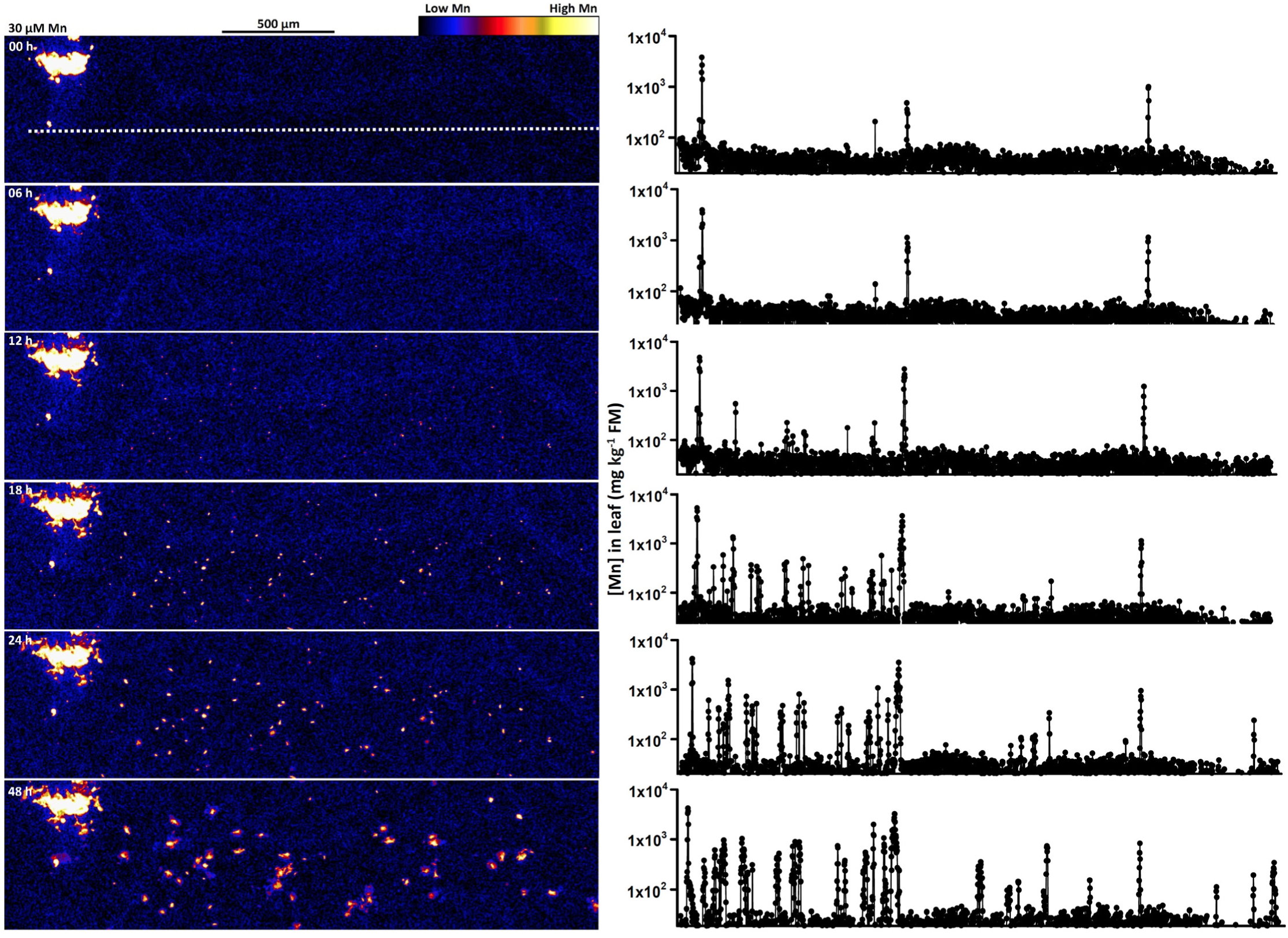
High-resolution μ-XRF scans and transects of Mn distribution and concentration (log10 scale) from 0 to 48 h after initial exposure to 30 μM Mn in a 2.6 mm × 0.6 mm section of a cowpea unifoliate leaf (Fig. 3). The horizontal dotted line at 0 h identifies the position of transects along which the concentration of Mn was determined.

As demonstrated by Scheckel *et al.* (2004), ImageJ analysis of the high-resolution data at 30 μM Mn (Fig. 4) extended the two-dimensional μ-XRF scans to a three-dimensional geometry providing visual images of both Mn distribution and concentration over the 48-h experimental period (Fig. 5). As with the two-dimensional images (Figs 3, 4), there were a few spots of high Mn at the start of the experiment with no discernable increase at 6 h. It appeared that the number and concentration of Mn in these spots started to increase at 12 h followed by a further increase at 18 h and marked increases at 24 h and 48 h.

**Fig. 5.**
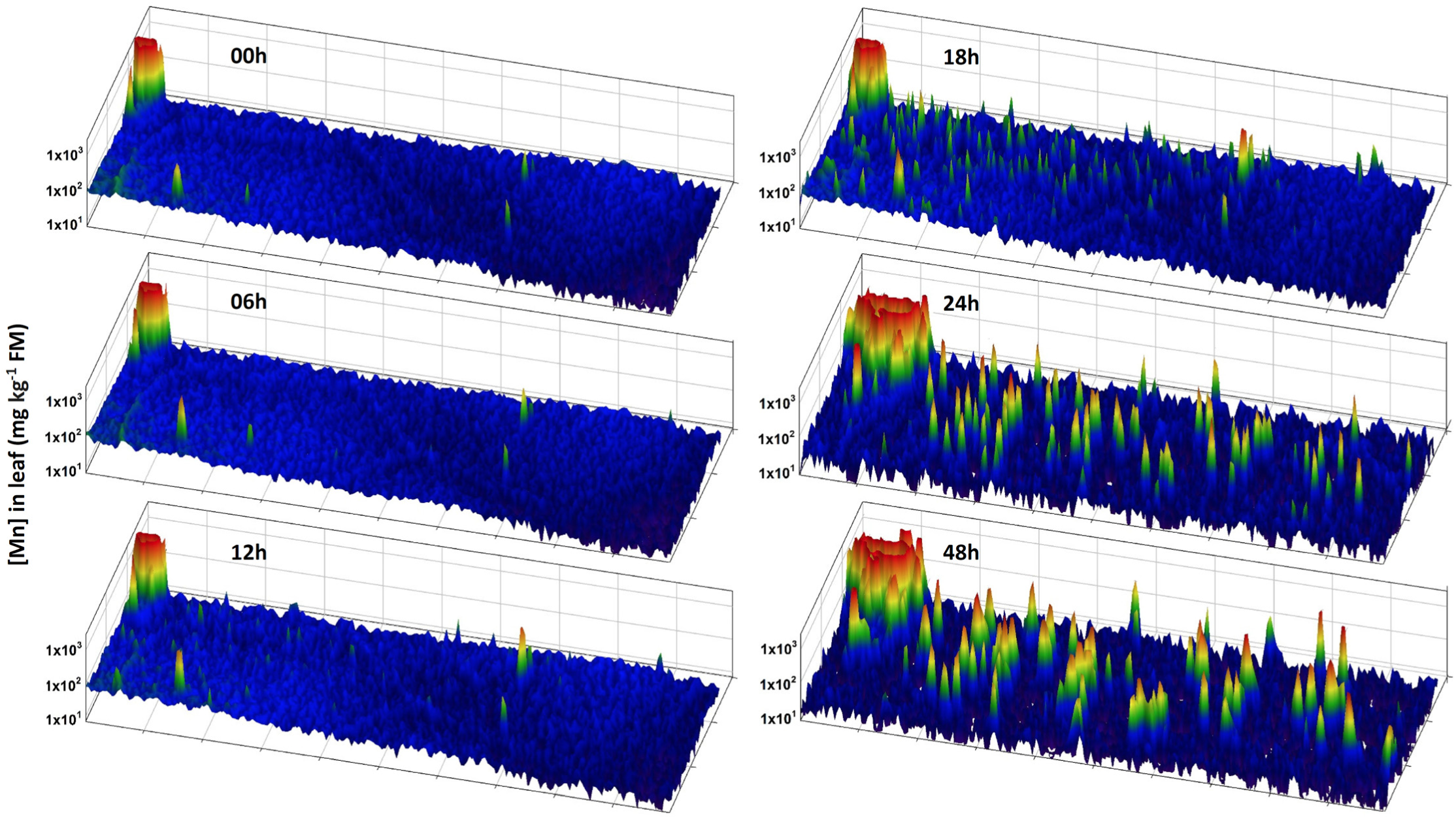
Three-dimensional representation of Mn distribution and concentration in unifoliate leaf sections of cowpea from 0 to 48 h after initial exposure to 30 μM Mn. The scanned area is 2.6 mm × 0.6 mm as shown by the red box in Fig. 3 and the high-resolution μ-XRF images in Fig. 4. The Mn concentration in the large high-Mn spot (back left) exceeded 10,000 mg kg^-1^ FM from 0 to 48 h but the scale of Mn in the unifoliate leaf has been limited to 1,000 mg kg^-1^ FM to show the change in high Mn spots that developed during the 48-h experimental period.

Using the detailed scan areas of 55.2 and 56.4 mm^2^ at 0.2 and 30 μM Mn (Fig. 3), we used ImageJ to calculate the number of localized Mn spots at 0 h was 3.6 and 12.1 mm^-2^ in the two leaves to be subjected to the 0.2 and 30 μM Mn treatments (Fig. 6a). The number of high-Mn spots appeared unchanged over the first 6 h in both treatments. There was a slight increase over 48 h of 4 mm^-2^ at 0.2 μM Mn compared to the large increase of 30 mm^-2^ at 30 μM Mn. The corresponding increases in the total area of localized high-Mn spots were < 0.001 and 0.04 mm^2^ mm^-2^ (Fig. 6b). Further analyses using ImageJ determined that most of the initial high Mn spots were < 320 μm^2^ in size (Fig. S2), largely remaining so over 48 h at 0.2 μM Mn. This contrasted to the changes in spot size in the 30 μM-Mn treatment in which the percentage of spots < 64 μm^2^ decreased over 48 h. Interestingly, the percentage of slightly larger spots of 64 to < 320 μm increased and then decreased and there was a concomitant increase in large spots at 48 h.

**Fig. 6.**
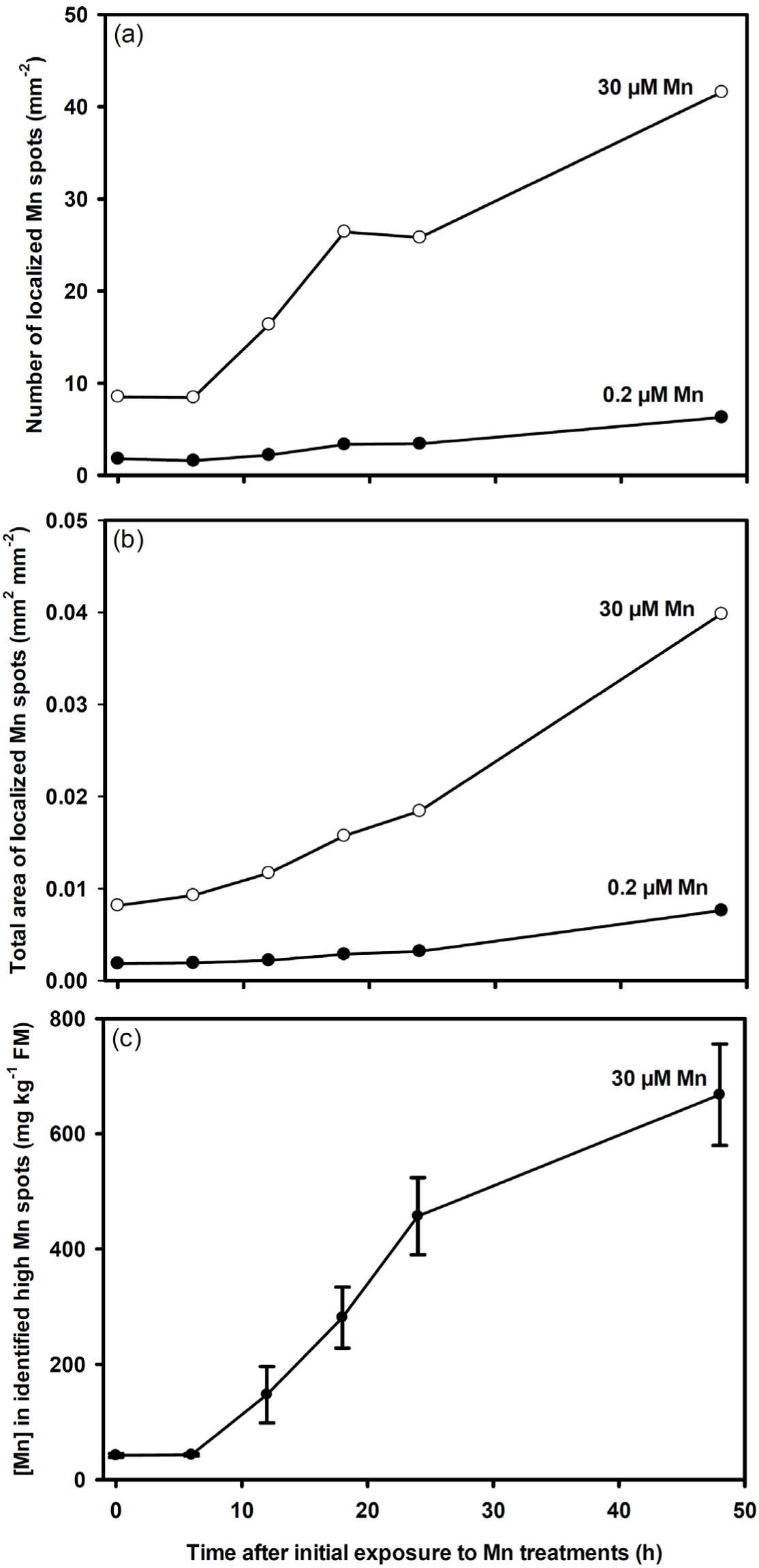
Accumulation of Mn in unifoliate leaf sections (Fig. 3) of cowpea from 0 to 48 h after initial exposure to 30 μM Mn in solution culture. (a,b) ImageJ determination of the number and area of localized spots of high Mn in detailed scans of leaf sections (Fig. 3). (c) GeoPIXE determination of the Mn concentration in identified high Mn spots (i.e. > 60 mg kg^-1^ FM) over 48 h in transects of the section of a cowpea unifoliate leaf at 30 μM Mn (Fig. 4).

With the μ-XRF scans providing quantitative, true-elemental data (Ryan & Jamieson, 1993; Ryan, 2000), GeoPIXE analysis of the detailed μ-XRF scan of leaf sections at 48 h (Fig. 3) determined mean values of 26 and 56 mg Mn kg^-1^ FM at 0.2 and 30 μM Mn. Analysis of the six transects resulted in a calculated mean Mn concentration that increased from 35 to 61 mg kg^-1^ FM over 48 h exposure to 30 μM Mn (Table S1). However, these transects indicated that the background Mn concentration was < 100 mg kg^-1^ FM other than in the relatively few instances in which the Mn concentration approached or exceeded 1,000 mg kg^-1^ FM. Separating the data into two classes, above and below an estimated background value of 60 mg kg^-1^ FM, permitted further examination of leaf Mn status. Pixel clusters with values < 60 mg Mn kg^-1^ FM had a mean concentration of 24 mg Mn kg^-1^ FM over 48 h, markedly lower than in those with values > 60 mg kg^-1^ FM that increased from 200 to 420 mg Mn kg^-1^ over 48 h (Table S1). Data from the 2.6 × 0.6 mm area used also to determine leaf Mn concentration increased from a mean of 69 to 170 mg kg^-1^ FM over 48 h (Table S1). These estimates differed somewhat from the the bulk Mn concentration in entire unifoliate leaves 48 h after imposing the high Mn treatment of 11 ± 1 and 47 ± 14 mg kg^-1^ FM at 0.2 and 30 μM Mn. (Corresponding values were 80 ± 6 and 360 ± 110 mg kg^-1^ DM.)

Finally, we utilized the μ-XRF analyses at 0 and 48 h to generate images for multiple elements within the energy range of the beamline. In this exercise, the incident X-rays of 12.9 keV and the Maia detector permitted investigation of the distributions and concentrations of elements between P and Zn. (Selection of a higher incident energy would also permit analyses of heavier elements.) The present study permitted investigation of six elements, K, Ca, Mn, Fe, Cu, and Zn, in unifoliate leaves that were above background concentrations (Fig. S3). At 0 h, there was relatively even distribution of K and Ca across the leaves of plants at 0.2 μM Mn, as was that of Cu though at a concentration close to the detection limit. In contrast, Mn and Fe accumulated in localized areas across the leaf, but these elements were not co-located. Localized areas of high Zn were present in and adjacent to the veins. The distributions of these elements were similar 48 h later at 0.2 μM Mn but the distribution of Mn differed between the 0.2 and 30 μM Mn treatments as is evident in Fig. 3.

## Discussion

The analysis of fresh tissues exacerbates the many challenges in μ-XRF scanning of low metal and metalloid concentrations in biological tissues (Lombi *et al.*, 2011a). These challenges arise from tissue hydration (often > 85 %) and the generally low concentrations of trace elements in tissues that result in the consequent long transit times required for analysis. A high (long) dwell increases the likelihood of radiation damage and tissue dehydration, thereby resulting in experimental artifacts that affect element distribution (Lombi *et al.*, 2011a). The benchtop (i.e. non-synchrotron based) μ-XRF method developed by Fittschen *et al.* (2017) ensured high sensitivity and low detection limits but required dwells of 1 s per 40 × 40 μm pixel that resulted in substantial radiation damage to a scanned *Arabidopsis thaliana* leaf. Scheckel *et al.* (2004) also noted that μ-XRF analysis using a 1-s dwell of a 6-μm pixel damaged leaves of living *Iberis intermedia* plants. As with these studies, our previous unpublished work showed radiation damage and leaf dehydration was evident in a soybean leaf subjected to multiple scanning (Fig. S1). Recent years, however, have seen major improvements in overcoming these problems in synchrotron-based studies. This has been important in the study of fresh, hydrated root (Kopittke *et al.*, 2011) and leaf tissues (Blamey *et al.*, 2015). Besides the high spatial resolution of 0.1 to 0.2 μm at the Australian Synchrotron XFM beamline (Paterson *et al.*, 2011), there are inherent advantages of the Maia detector and Rascan in decreasing dwell. This was evident in the preliminary experiment of the present study in which there was no visible X-ray damage to a unifoliate cowpea leaf after two survey scans and four detailed scans with a combined period of *c*. 2 h (Fig. 1). The recently updated raster scanning adds further advantages by decreasing the damage to sensitive plant tissues.

Progress in μ-XRF analysis of hydrated plant tissues has been extended to leaf tissues of living plants. For example, by growing the hyperaccumulator *I. intermedia* plants at elevated Tl in soil, Scheckel *et al.* (2004) determined that Tl accumulated mostly in the vascular tissues. Fittschen *et al.* (2017) constructed a benchtop μ-XRF unit to examine differences in the distributions of a number of nutrients in the vascular system, mesophyll, and trichomes of leaves of living *A. thaliana* plants. These studies, however, did not investigate time-resolved changes in elemental distribution and concentration and we were not able to find instances in the literature of sequential μ-XRF analysis of living plant tissues.

In the present study, μ-XRF detailed scans using a pixel size of 4 × 4 μm enabled production of high-quality images that permitted direct visual comparisons between the effects of 0.2 and 30 μM Mn (Fig. 3) and enabled high resolution comparisons of Mn accumulation of 30 μM Mn over 48 h (Fig. 4). A pixel size of 1 × 1 μm allows even better resolution where appropriate (Blamey *et al.*, 2015). As demonstrated by Scheckel *et al.* (2004) and in the present study (Fig. 5), three-dimensional representations of the two-dimensional scans provide further visualization of elemental distribution and concentration.

Until recently, quantitative elemental analyses have focused on bulk tissues, from whole plants to separate analyses of roots, shoots, stems, and leaves (Salt *et al.*, 2008) but technological developments in synchrotron-based μ-XRF, amongst other techniques, provide greater spatial resolution of the ionome (Punshon *et al.*, 2009; van der Ent *et al.*, 2017). However, van der Ent *et al.* (2017) urged caution in the use of μ-XRF in foliar studies using living plants because measurements arise from different depths, from different cell types (e.g., epidermis, palisade, mesophyll, and vasculature), and from continued plant metabolism. The first two of these potential difficulties may be addressed by a combination of analytical techniques as shown by Blamey *et al.* (2017) who used NanoSIMS to confirm the suspected accumulation of Mn in the apoplast using μ-XRF by analysis. The present study has also shown that accommodation of continued plant metabolism is possible, albeit in only one instance of Mn accumulation. High Mn accumulation in discreet locations is of additional benefit by overcoming the overall low concentrations of many trace elements.

It has been recognized for many years that symptoms of Mn toxicity precede a decrease in plant growth (Foy *et al.*, 1978; Weil *et al.*, 1997), information extended by Fernando *et al.* (2016) with Mn accumulation in leaves of wheat (*Triticum aestivum*) plants well before the appearance of visible symptoms. This was evident in leaves of cowpea exposed to a non-toxic concentration of 0.2 μM Mn (Figs 1, 3, 4, 5). Localized spots of high Mn preceded visible symptoms associated with Mn toxicity in the present study also with new high Mn spots first visible after 12 h growth at 30 μM Mn (Figs 4, 5) but visible dark spots appeared only after 48 h (Fig. 2c). However, the mechanism is unknown whereby high Mn accumulates in specific localized areas not visibly associated with leaf anatomy (e.g., close to or distant from veins). There is similar uncertainty as to why an overall increase in Mn in leaf tissues at 30 μM Mn (Figs 4 and 5) results in localized increases in Mn but no general increase in background Mn concentration over time.

From an ionomics viewpoint (Salt *et al.*, 2008), the present study addressed various microscopic measures of Mn concentration in leaf tissues based on the true-elemental data of the μ-XRF scans. After 48 h at 0.2 μM Mn, these analyses ranged from 24 to 35 mg Mn kg^-1^ FM compared to the unifoliate leaf bulk analysis of 11 mg Mn kg^-1^ FM. The corresponding comparison was 56 to 169 mg kg^-1^ versus a bulk leaf tissue concentration of 47 mg Mn kg^-1^ FM at 30 μM Mn. While of the same order of magnitude, these findings suggest that further research is required in the sampling for quantitative determination of elemental concentrations in leaf tissues. Importantly, the multi-element analyses undertaken in the present study would provide important information on interactions between the various elements. An example relevant to the present study might use detailed μ-XRF studies to examine the kinetics of changes in Mn distribution (Figs 4, 5) with those of Ca (Fig. S3) given the role of Ca in callose formation associated with Mn toxicity (Wissemeier & Horst, 1987). Additional studies may also provide critical information as to the underlying mechanisms whereby Mn is toxic by determining changes in Ca and Mn accumulation in the apoplast and by determining how Mn accumulates along with Ca in vacuoles of some plant species but not in others. Finally, there is particular promise in studies such as that by Takahashi *et al*. (2009) in combining information on the activities of metal transporter genes with multi-element μ-XRF analysis of changes in the distribution of Fe, Mn, Cu, and Zn in germinating rice (*Oryza sativa*) seeds.

## Conclusion

Development of a synchrotron-based method has shown the feasibility of using μ-XRF analysis to determine the distribution and concentration of Mn in living unifoliate leaves of cowpea exposed to 0.2 and 30 μM Mn in solution culture. This method may be adapted to examine the accumulation and distribution of other elements and plant species over a range in exposure times. Besides investigations of elemental toxicities, other examples may include determining the dynamics of nutrients upon addition to a nutrient-deficient plant, the effects of hypoxia on nutrient status, and establishing the relationship between transpiration and nutrient accumulation in leaf tissues.

The results and implications of the present study advance the value of ionomics, the inorganic component of the metabolome, through quantifying changes in the microscopic distribution and concentration of elements *in vivo* (Salt *et al.*, 2008). Though focusing the present study on the kinetics of Mn accumulation in cowpea unifoliate leaves, μ-XRF analyses may also provide multi-element information in plant tissues, meeting an aim of ionomics to define the functional state of an organism driven by genetic, developmental, biotic, or abiotic influences.

## Acknowledgements

F.P.C.B. acknowledges assistance from the Australian Government Research Training Program and P.M.K. acknowledges receipt of an Australian Research Council (ARC) Future Fellowship (FT120100277). This research was undertaken on the XFM beamline (Project AS153/XFM/11040) at the Australian Synchrotron, part of the Australian Nuclear Science and Technology Organization (ANSTO). We thank Dr Chris Ryan (CSIRO), Dr Martin de Jonge, and Dr Daryl Howard (Australian Synchrotron) for assistance with synchrotron-based techniques.

## Author contributions

F.P.C.B. conceived the research program that was developed in collaboration with D.J.P., A.W., N.W.M., and P.M.K.; A.W. and N.A. developed the requisite technology; F.P.C.B., M.C., and C.T. conducted the plant growth experiments; F.P.C.B., D.J.P., B.A.M., M.C., and P.M.K. conducted the μ-XRF analyses at the Australian Synchrotron; F.P.C.B and P.M.K. analyzed the data; and F.P.C.B., D.J.P, and P.M.K. wrote the first draft of the article to which all authors contributed.

